# Paediatric MRI: High-Resolution *in vivo* 3T Methods

**DOI:** 10.1101/2025.10.30.685686

**Authors:** Brooklyn R. Wright, Mark M. Schira, George Paxinos, Mustafa S. Kassem

## Abstract

Magnetic Resonance Imaging (MRI) is a powerful tool for investigating the brain *in vivo* but is limited by image resolution and scan artefact. In paediatric research, these limitations are compounded by reduced participant compliance, which in turn necessitates shorter scan times to minimise motion artefacts, resulting in less than satisfactory image resolution. We report here methods for MRI acquisition which afford high-resolution, low-noise, paediatric brain data in under nine-minute scans, and novel post-processing with our code freely available at https://osf.io/ckh5t/. Whole-brain *in vivo* single-participant images were generated at 0.5mm isotropic resolution. This resolution permitted accurate delineation of fine structures, including the hippocampal subfields. The ability to resolve intricate structures in paediatric data provides a tool for studying brain development and its disruption in neurologic and psychiatric disorders.

## 1. Introduction

While almost all neurons are present at birth, the brain undergoes a period of accelerated growth in the first five years (Van Dyck & Morrow., 2017). Brain development is a highly complex process, characterised by the remodelling of neural circuits, something that persists into the early twenties, with grey matter density increasing until age thirty (Ball et al., 2021; Sowell et al., 1999; 2003). Despite advances in MRI technology, data resolution, and image clarity, there are no high-quality paediatric MRI data; hindering capacity to investigate vital changes as the human brain matures. A significant challenge faced during paediatric scanning is participant compliance, children finding it difficult to remain sufficiently still in the MRI. This increases the likelihood of motion artifacts, something that compromises data quality (Greene et al., 2016; Makowski et al., 2019). Elimination of the artifacts is made more difficult by high resolution imaging which often requires long scan times (>20-minute), where head motion, even as subtle as swallowing or twitching, affects scan quality (Schira et al., 2023).

Because of motion artifacts, collecting child MRI at high resolution is often considered impossible without the use of sedation (Greene et al., 2016). Sedatives introduce additional risks and, therefore, are not favoured (Arlachov & Ganatra, 2012). As a result, there is a shortage of high-quality, normative paediatric MRI data with sufficient resolution to reliably delineate either cortical or subcortical structures. Yet, high-resolution imaging is essential for visualising anatomical boundaries accurately. For example, the hippocampus—a structure critical for memory—is composed of layered grey and white matter that can only be resolved with sub-millimetre resolution imaging. The unavailability of child MRI at high resolution limits research and clinical interpretations. For example, in the assessment of epilepsy and neurodevelopmental disorders such as attention deficit hyperactivity disorder (ADHD), high-resolution imaging can indicate morphological changes typically present earlier than clinical symptoms (Bedford et al., 2025; Gilbert et al., 2001; Priol et al., 2021; Zhao et al., 2022).

Without the detail provided by high resolution data from canonical paediatric populations, understanding the trajectory of human brain development relies heavily on the extrapolation of adult data or animal models (Zhao & Bhattacharyya, 2018). Both approaches have inherent limitations, restricting our ability to form a comprehensive understanding of (a) healthy brain development, and (b) neurological and psychological disorders in children. Critically, using adult brain anatomy as a framework to study developmental change presupposes what it aims to discover. This circularity is concerning given that grey and white matter structures undergo dynamic, region-specific and sex-specific changes during development. Modern delineation techniques depend heavily on spatial priors — when structural boundaries are unclear, they are inferred based on adult anatomy. This dependency is the consequence of a fundamental gap: the absence of child-specific, high-resolution anatomical data. Hence, we advocate the need for accessible methods to obtain high quality paediatric data with the goal of understanding the developing brain in both research and clinical practice.

Advancements in MRI technology, including the development of high-field systems like 7 Tesla (7T) and 9.4 Tesla (9.4T), along with enhanced acquisition protocols, have significantly improved imaging resolution (Isaacs et al., 2020). For example, the Schira et al. (2023) *Human Brain Atlas Project* achieved low-noise, whole-brain *in-vivo* structural MRI for the adult at a spatial resolution of 0.25mm isotropic, 64 times finer than the conventional 1mm³. Schira et al. (2023) obtained structural detail approaching histology by implementing intensive scanning protocols requiring adult participants to undergo approximately 20 sessions and combining this with advanced post-processing and averaging techniques. Despite these breakthroughs, 7T systems and high-resolution protocols remain largely impractical for routine research and clinical applications due to their limited accessibility and higher costs compared to ubiquitous 3 Tesla (3T) systems (Beisteiner et al., 2011; Boubela et al., 2014). Consequently, there is a need for optimised MRI acquisition and processing techniques that utilise 3T systems to deliver image quality and resolution typically associated to 7T systems (Bahrami et al., 2016).

Recent work aiming for high quality data of individual adult subjects has adopted data processing pipelines initially developed for multi-subject template generation (Lüsebrink et al., 2017; Schira 2023). Template generation is a neuroimaging software method used to create standardised reference targets by morphing and averaging multiple MRI datasets (Pai et al., 2020). When applied to single participant data, that is multiple scans of the same brain, the method of template generation can be repurposed to significantly decrease the signal-to-noise ratio (SNR) improving image quality and resolution, a technique known as ‘super-sampling’ (Shilling et al., 2008; Van Reeth et al., 2012). Super-sampling allows researchers to push for higher resolution images from MR scanning. Here we demonstrate these techniques on paediatric data and suggest their use in clinical applications.

We used refined data processing procedures from Schira et al. (2023), though modified for 3T scanning and shorter acquisition times (acknowledging the specific needs of paediatric cohorts), and achieved increased resolution and clarity compared to conventional 3T MRI data. We first demonstrated and tested these adapted methods on adult participants to assess their feasibility and utility before applying them to our child participants. Adult participants underwent four MRI sessions to render multiple high-resolution T1-weighted (T1-w), T2-weighted (T2-w), and diffusion weighted imaging (DWI) scans, while child participants underwent two acquisition sessions rendering solely structural data (T1-w and T2-w scans).

## 2. Methods

### 2.1 Participants

Three healthy adult participants and two healthy children with no known neurologic or psychiatric pathology participated in this study. The adult participant group consists of two female and one male volunteer (aged from 21, 26 and 39, respectively). The child participant group consisted of two female volunteers (aged 9 and 12). Participants were screened by an independent radiographer to confirm they were suitable for MR scanning, e.g., no metal implants. Participants were given detailed information and consent forms, followed by an in-person consultation to verbally explain the study requirements and potential risks. Informed consent was then collected. For child participants, consent was obtained from both the parent/guardian and the child. Children received age-appropriate information and consent forms, which included pictures and clear, accessible language. To improve participant comfort, participants were given the choice to watch videos or listen to music during scans, with TV show ‘Bluey’ being a popular choice in the child participant group.

### 2.2 Scanning Acquisition Parameters

Scans were acquired using a whole-body Philips Ingenia CX 3T MRI, narrow bore (60cm) system. The number of repeated acquisitions varied across participants. For the female adult participants, six T1-w scans and five T2-w scans were collected. The male adult participant underwent five T1-w and four T2-w scans. A significant portion of the scanning time for adult participants was dedicated to acquiring DWI data. While the development of the DWI protocol was successful for adult scanning, the high-resolution DWI protocols were deemed too long to be suitable for child participants. These DWI protocols will be the subject of a different report. For the child participants, four T1-w scans and four T2-w scans were obtained because analysis of the adult data suggested 4 scans are sufficient to achieve the desired quality improvements.

T1-weighted images were acquired at 0.65 mm isotropic resolution in the sagittal plane using a 3D-MPRAGE sequence (TR = 2000 ms, TE = 3.59 ms, TFE factor = 157, flip angle = 8°). Parallel imaging was applied with a CSENSE reduction factor of 2, using an acquisition matrix of 368 × 368 mm and a bandwidth of 216 Hz/Px. The total scan time was 8 minutes 45 seconds.

T2-weighted images were also acquired at 0.65 mm isotropic resolution in the sagittal plane using a 3D Turbo Spin Echo sequence (BrainVIEW T2; TR = 3000 ms, TE = 270.79 ms, flip angle = 90°). Parallel imaging was applied with a CSENSE reduction factor of 8, using an acquisition matrix of 368 × 370 mm and a bandwidth of 828 Hz/Px. Each T2-weighted scan required 8 minutes 24 seconds.

### 2.3 T1-w and T2-w Image Pre-Processing

All acquisition scans underwent identical pre-processing to de-identify the data and improve image quality before being run through template processing. Pre-processing included the generation of brain-masks for all T1-w and T2-w images using the HD-Bet tool to identify and separate cortical tissue from skull and non-cortical tissue (Isensee et al., 2019). The Advanced Normalisation Tools (ANTs) ’Apply Transforms’ filter was used to realign the transformation matrix of the mask to its corresponding MR image before subsequent pre-processing steps (Avants et al., 2011). This was followed by ANTs ‘N4BiasFieldCorrection’ to correct variations in luminance intensity across the image. Following luminance inhomogeneity correction, the brain masks were dilated by a generous border of 12 voxels. Expanding the mask boundaries facilitated a subsequent skull strip of the MR images, that removed excess non-brain data from the scan whilst retaining a buffer zone of dura and skull to diffuse the effects of warping at the image boundary during the subsequent template registration process. The luminance-inhomogeneity corrected images were skull-striped using ANTs ‘ImageMath’ tool (Avants et al., 2011).

### 2.4 Template Generation

A first pass non-linear aligned average was generated for each of the T1-w and T2-w scans for all participants. To achieve this, images were first collated in a stack and aligned into the same template space. These aligned images were then averaged using the ‘mrregister’ command from the Mrtrix 3.0 suite (Tournier et al., 2019). The affine-nonlinear option was specified as well as a scale of 1.5 to facilitate resolution super-sampling, producing an output image with an increased resolution of 0.4mm^3^ from the acquisition 0.65mm^3^. These averaged images were rescaled to our final resolution of 0.5mm^3^ isotropic using the Mrtrix 3.0 mrgrid tool. This was done to maintain improved image resolution and optimal template registration refinement, whilst ensuring the final templates retain a resolution compatible with the post-processing and reconstruction software. Finally, these images were re-oriented into the Anterior Commissure (AC) – Posterior Commissure (PC) ‘ACPC’ space. This is a standardised coordinate system that uses the AC and PC as references to make the horizontal plane in histology and MRI research (Bhanu Prakash et al., 2006).

The purpose of this initial template is to establish a standardised target space and grid for the final dataset, ensuring the brain image was in ACPC orientation, had an exact voxel size of 0.5mm^3^ and transformation matrices that defined the resolution and the centre of the coordinate system to be at the AC. A key advantage of this standardisation is that it also enables seamless integration into Python and other programming environments allowing slices to be displayed in the ACPC orientation without requiring further orientation transformations which risk image quality reduction. Since this template served as the starting point for subsequent averaging, mild blurring introduced during this initial re-slicing process had no impact on the quality of the final image.

Final super-sampled average images for each contrast and participant were created using ANTs ’antsMultivariateTemplateConstruction2’ script (Avants et al., 2011. The script requires a starting template, and for this served the initial single-participant average template described above. The initial template is iteratively refined and replaced three times through the non-linear alignment of the template’s constituent acquisition scans. During this template generation process the following parameters were specified: (a) the scans were aligned using a cross-correlation similarity metric (-s flag), (b) a Greedy SyN transformation model (-t flag) was used to implement non-linear registration, (c) the maximum number of step sizes (-m flag) was set to 20×15×5 and (d) a gradient (-g flag) of 0.1 was employed. The initial template image was specified as the starting point of normalisation using the -z flag. These techniques were modified based on the Schira et al. (2023) methods for 7T ultra-high resolution template generation, which themselves are based on Lüsebrink (2021).

It is important to note that due to the continued development of the ANTs tools, different version and script iterations of ANTs software result in some output variability. Thus, for reproducibility, specific software modules must be employed. For the adult data, version v2.3.5.post95-g6f07ac5 (Feb 15. 2022) was employed. For compatibility and reproducibility reasons, we switched to a Neurodesk module for the child datasets (module: ants/2.3.5; Version: 2.3.5.dev208-g6f137, Dec. 10. 2021). All processing methods and scripts can be found at (OSF: https://osf.io/ckh5t/) and are compatible with Neurodesk which allowed for specification of software versions (Renton et al., 2024).

## 3. Results

Our results demonstrate the anatomical clarity achievable through combining sub-millimetre time-efficient 3T protocols with refined post-processing super-sampled averaging methods. Figure 1 displays the final whole-brain 0.5mm isotropic T1-w and T2-w templates for our 9-year-old and 12-year-old child participants, and the 21-year-old adult participant. Our template generation process, when applied to single participant datasets, permits super-sampling (Schira et al. 2023). The final image resolution of 0.5mm^3^ isotropic surpasses the acquisition resolution of 0.65mm^3^ isotropic. At first glance, a voxel reduction of 0.15 mm³, from 0.65mm^3^ to 0.5mm^3^, appears modest; however, it represents a 54% reduction in voxel volume and a 2.2× increase in spatial sampling density. Furthermore, achieving 0.5mm^3^ resolution is an eightfold increase in voxel count compared to the conventional 1mm^3^ resolution. This substantial gain occurs because reducing the voxel size in each dimension (x, y, and z) leads to a cubic increase in the total number of voxels, and by virtue of this, image information (see Fig. 4F).

**Fig. 1.**
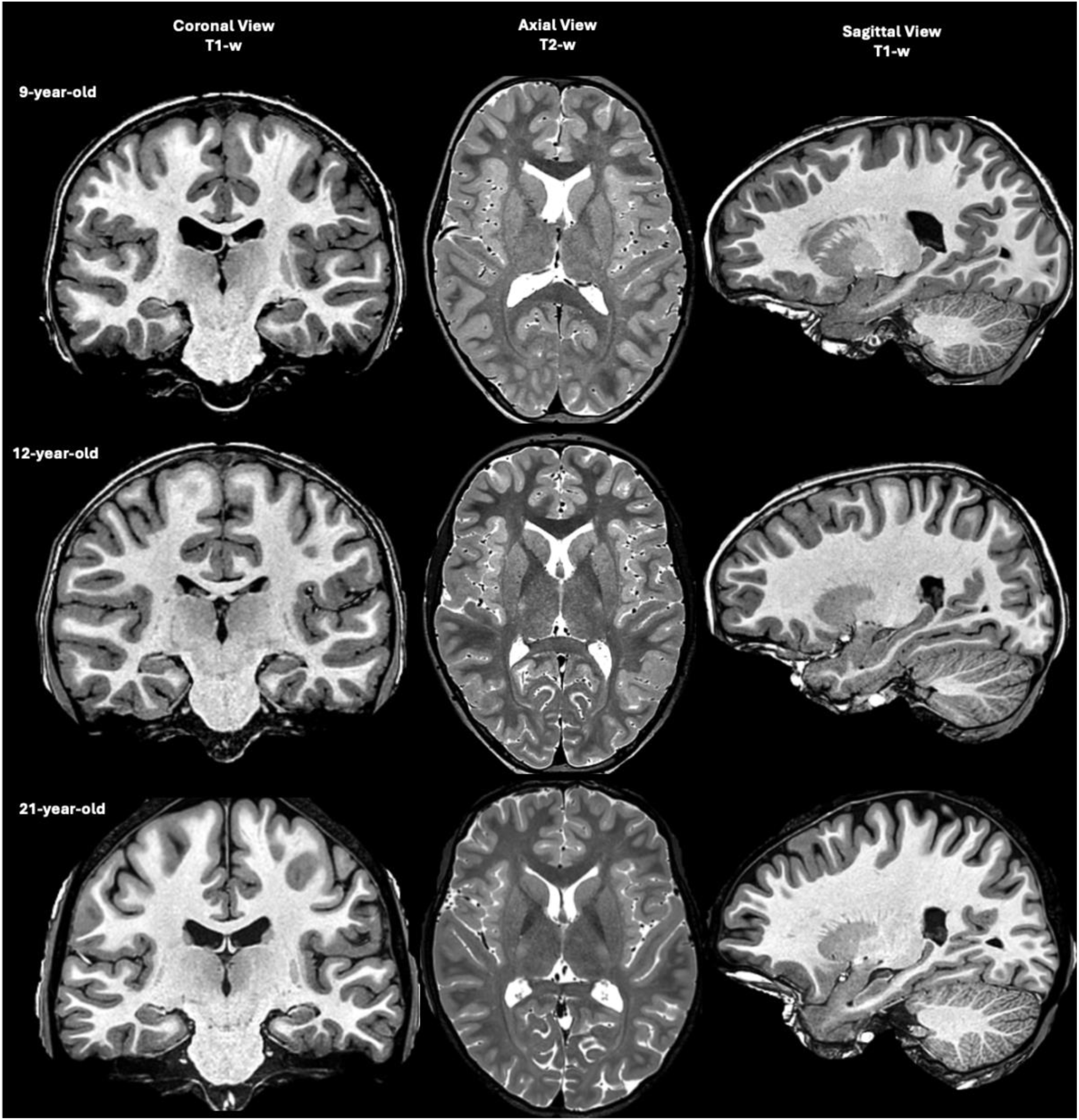
Whole Brain Final Images at 0.5mm Isotropic in Three Cardinal Planes. Final whole-brain images for each participant displayed across three anatomical planes. The top row shows images from the 9-year-old child, the middle row from the 12-year-old child, and the bottom row from the 21-year-old adult. Within each row, the coronal (left, T1-w), axial (middle, T2-w), and sagittal (right, T1-w) views are presented.

To assess the stability of the MRI acquisition parameters and quantify signal variation across repeated whole-brain measurements, a voxel wise relative standard deviation (RSD) analysis was performed (Fig. 2). For each paired voxel, the standard deviation (SD) of luminance intensity values across the repeated acquisitions was calculated. This was normalised by dividing this SD by the mean signal intensity for that voxel and converted into percentage. To improve robustness a gaussian filter was applied to smooth the RSD values (but not the original images). The highest levels of signal variability were observed in cerebrospinal fluid (CSF) regions, which may reflect physiological motion, as well as partial voluming effects (Feinberg & Mark, 1987). To quantify signal variability in grey matter specifically, a FSL Fast segmentation was performed to mask the grey matter voxels (Jenkinson et al., 2021). The mean RSD across grey matter was 6.9%, this in turn allowed a simple power analysis estimating the expected standard error for each voxel when averaging multiple acquisitions (Fig. 3)

**Fig. 2.**
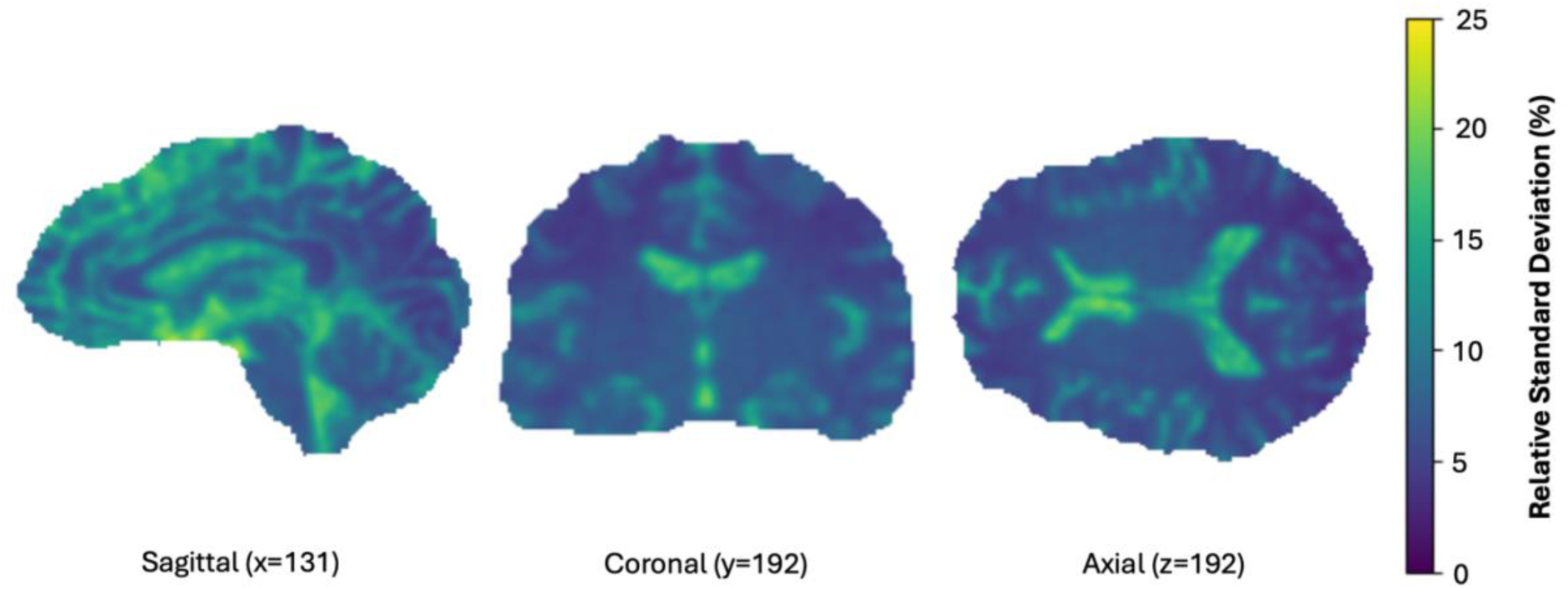
Relative Standard Deviation of Voxel Luminance Values Across Repeated Acquisition Scans. Relative standard deviation (RSD) map derived from six repeated T1-w acquisitions of our 26-year-old adult participant. Images show the average voxel wise RSD overlaid on anatomical space, in the sagittal (left), coronal (middle), axial (right) views.

**Fig. 3.**
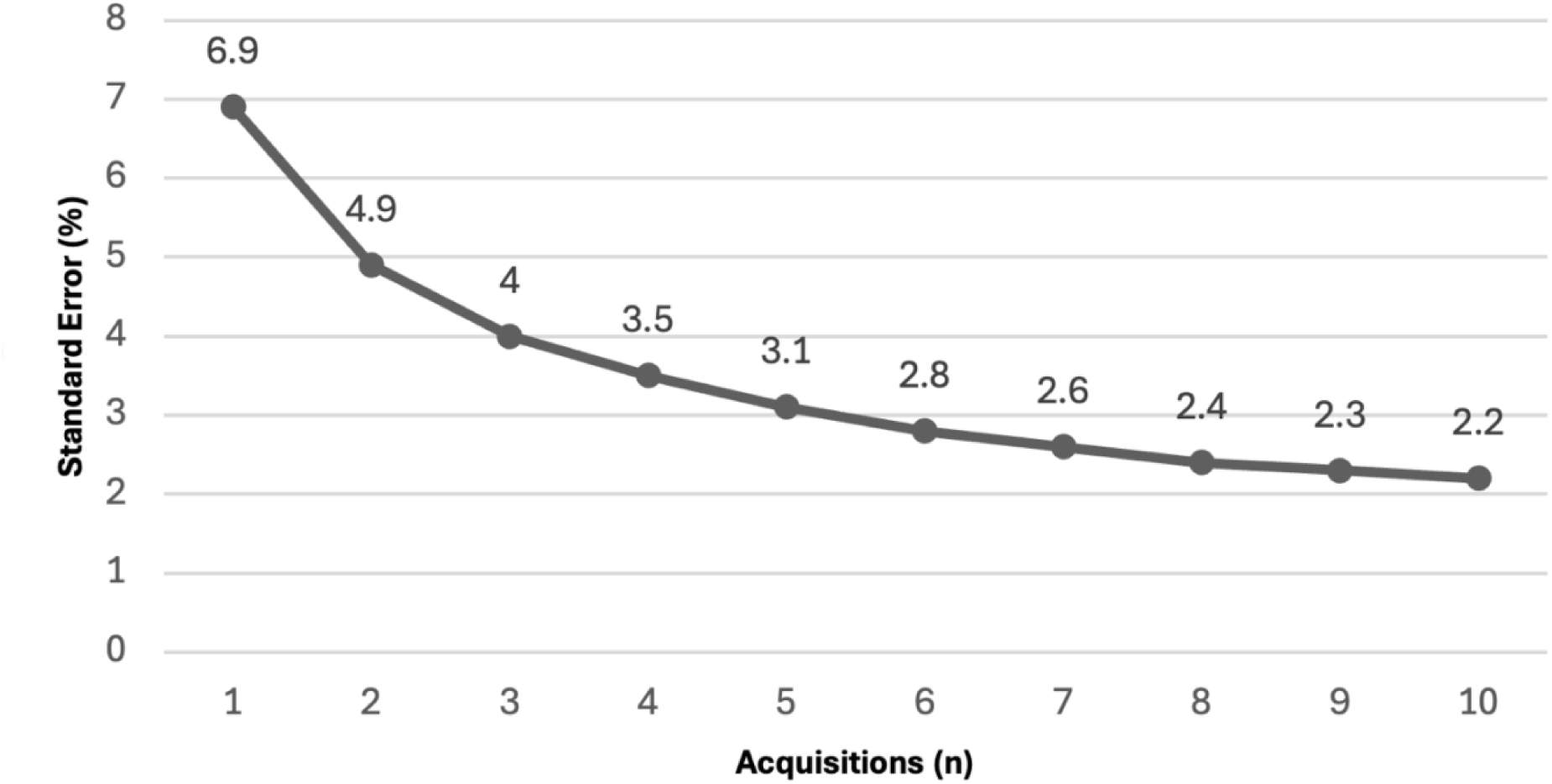
Reduction of Relative Standard Error of Voxel Luminance Values Across Repeated Scans. The reduction in relative standard error (RSE) of voxel luminance value as a function of the number of repeated acquisitions averaged.

**Fig. 4.**
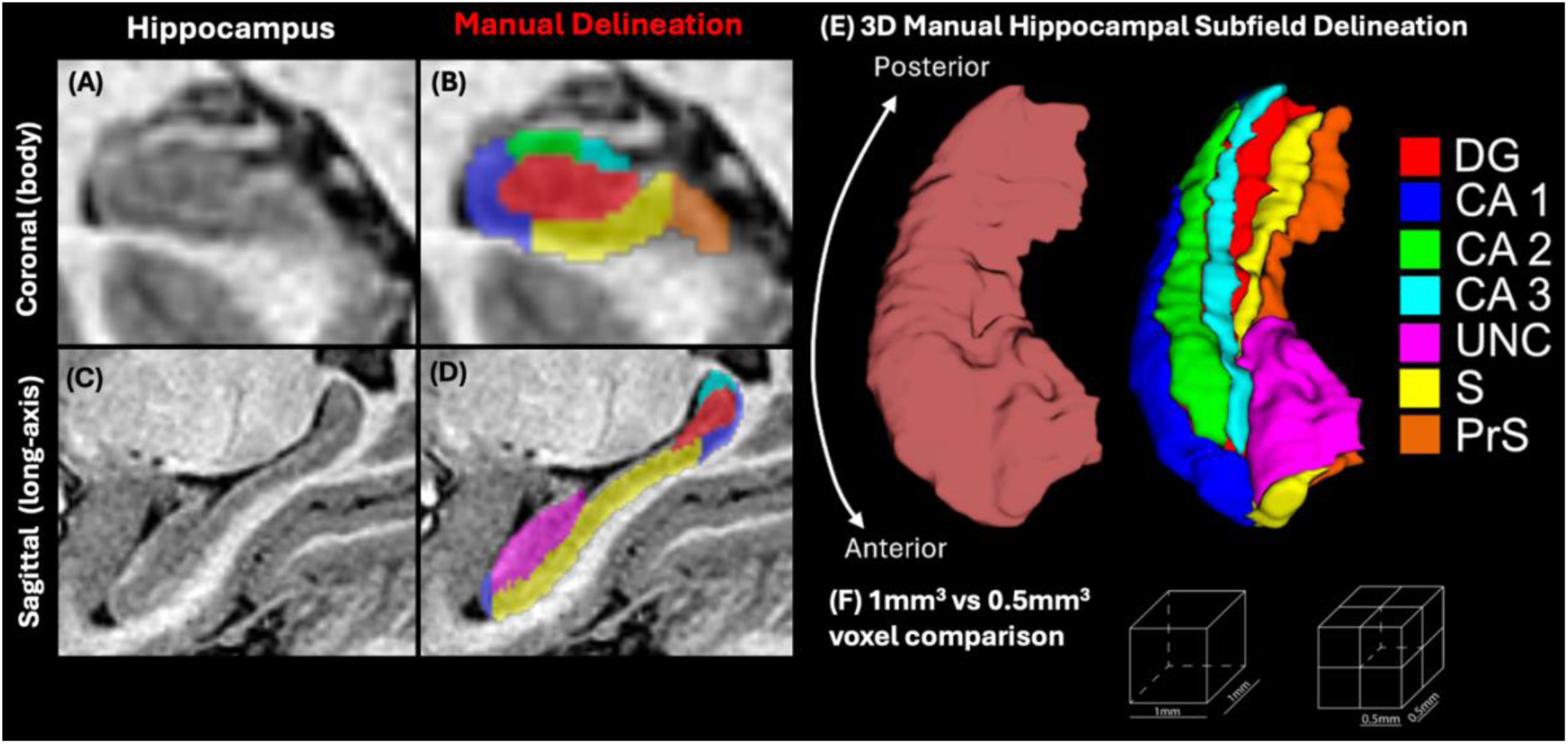
Manual Hippocampal Delineation of Paediatric (9-year-old) MRI. (A) T1-w 0.5mm³ MRI displaying the body of the hippocampus in the coronal plane. (B) The same coronal slice of the hippocampus with delineations overlaid. (C) T1-w 0.5 mm³ MRI of the hippocampus (long-axis) in the sagittal plane. (D) The same slice with delineation overlaid (E) Three-dimensional reconstruction of hippocampal delineations: the left panel shows the whole hippocampus, and the right panel displays the delineated subfields. (F) An illustration of the resolution gain when a single 1 mm³ voxel is split into eight 0.5 mm³.

Given the repeated acquisitions provide repeated measures of the same volume, we estimated the reduction in relative standard error (RSE) delivered by averaging multiple scans. Assuming a RSD of 6.9% in grey matter, the standard error for voxel luminance measurements decreases to 3.5% with four repeated scans and to 2.8% with six scans (Fig. 3). As the reduction in RSE becomes asymptotic with increasing scan number, the marginal gain in measurement precision from six acquisitions versus four was approximately 0.7%. This small improvement was deemed insufficient to justify additional scan time in paediatric participants. Therefore, the number of structural scans for child participants was reduced to four for each contrast (T1-w and T2-w), leading to a 33% reduction in acquisition time in the paediatric protocol.

To assess whether our MRI protocols produced images with satisfactory structural information, we delineated a structure with high complexity, i.e., the hippocampus. Since there is currently no hippocampal delineation protocol for children, this delineation was guided by an illustrated delineation protocol for 3T MRI for the adult (Dalton et al., 2017). Our delineation was conducted along the long-axis of the hippocampus in the coronal plane working from anterior to posterior. Our delineation of CA2 and CA3 subfields relied on geometric rules guided by histological knowledge which outlines CA2 as approximately one third the size of CA3 (Diers et al., 2023; Insausti & Amaral., 2012; Insausti et al., 2023; Mai et al., 2016). The clearly visible stratum radiatum and the stratum lacunosum-moleculare (SRLM) permitted delineation of the hippocampus into seven of its constituent subfields: the dentate gyrus, CA1-3, the uncus (unc), the subiculum (s) and the presubiculum (PrS).

As MRI resolution increases, the clarity of *in vivo* visualisations of neuroanatomy approach *ex-vivo* histology. Figure 5 illustrates improvements in structural definition between typical 1mm isotropic 3T MRI data from (Cole et al., 2025; Fig. 5A), to our single-scan acquisition data (Fig. 5B) and our final (Fig. 5C) image data. It is noteworthy that in typical MRI data (Fig. 5A), be it child or adult, the ‘hook’ of the PC is not visible, whereas it becomes clearly visible in our high resolution images. Improvements in structural clarity are most pronounced in the final image data (Fig. 5C) which permits the clearest visualisation of the hook and individual branches of cerebellar arbour vitae. The PC hook is essential for determining the ACPC plane because the PC end of the plane must pass through the centre of the PC (Bhanu Prakash et al., 2006). This plane is necessary for histological and MRI investigations and permits reliable comparison across research.

**Fig. 5.**
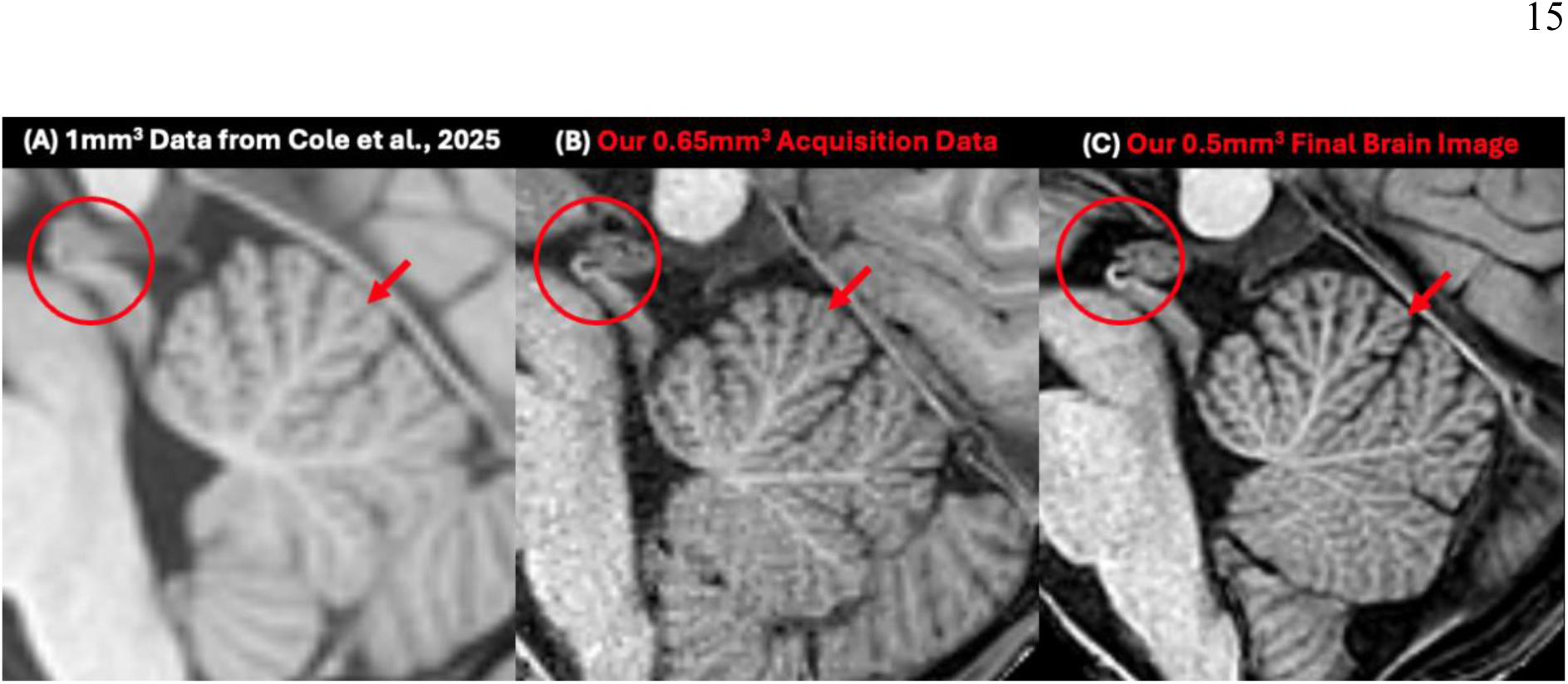
Comparison between typical MRI and our data for a 9-year-old. (A) Typical 3T data at 1mm^3^ isotropic resolution. The hook of the posterior commissure (red circles) and the arbor vitae of the cerebellum (red arrows) are not resolved (MRI data acquired from Cole et al., 2025). (B) Minimally processed data from a single acquisition at 0.65mm^3^ from a 9-year-old in our study; note the clearer outline of the hook of the posterior commissure, and (C) final super-sampled whole brain data at 0.5mm^3^ from our child dataset displaying further improved visibility of the posterior commissure and cerebellar microstructure.

## 4. Discussion

The MRI acquisition and processing methods presented herein provide high-resolution, high-contrast adult and paediatric brain data, attained within short scan times and without sedation. This demonstrates the feasibility to collect paediatric MRI data with significantly better quality than previously achieved. This was accomplished by splitting MRI acquisition across multiple time-efficient, sub-millimetre 3T scans for each participant which were then averaged and super-sampled. The MRI parameters in this study permitted the high-resolution acquisition (0.65 mm³) of T1w and T2w scans. Additionally, template generation software was employed to super-sample the data resulting in a final image resolution of 0.5 mm³, surpassing that of the original acquisition resolution by 0.15mm^3^. By adapting template generation software and using non-linear alignment, our methodology: (1) reduced artefact caused by head movement, the principal hurdle with child participants, (2) improved SNR through averaging multiple scans of the same participant, and (3) preserved fine details often not captured in single acquisitions (Shaw et al., 2019; Shilling et al., 2008; Van Reeth et al., 2012). The final brain images offer improved spatial resolution and microstructural clarity that surpasses typical datasets, including high quality datasets on adult participants, such as those from the Human Connectome Project (Somerville et al., 2018) and the highest-resolution paediatric brain atlas (Molfese et al., 2021). Critically, this afforded us greater accuracy in the identification of fine brain structures *in vivo*.

The gain in resolution not only enhances the visibility of fine structural details but also strengthens confidence in the delineation of brain structures. MR images are a reconstruction of luminance values derived from signal measurements. These measurements and their accuracy are influenced by the physics of MRI and its inherent limitations, such as field inhomogeneities, meaning the resultant image is always an approximation rather than a perfect capture. In line with the Nyquist–Shannon Sampling Theorem, which states that sampling must occur at least twice the highest frequency to prevent aliasing, higher spatial resolution is essential for improving the accuracy of boundary delineation and volumetric precision (Hu et al., 2014). Our protocol, which delivers 0.5 mm isotropic resolution, supports precise morphological and volumetric investigations of complex neuroanatomical structures, enabling visualisation of fine anatomical features that are undetectable in supra-millimetre (>1 mm³) 3T data, such as the posterior commissure and hippocampal subfield boundaries.

Our final whole brain images permitted the delineation of hippocampal subfields in child data. Hippocampal investigations are challenging even in adults, a result of both the limited visibility of the structure at low resolution, and scan artefacts that blur the SRLM band, a landmark necessary for delineation (Canada et al., 2024). While it is not possible to resolve layers that compose the SRLM in MRI, together they form a band consisting mostly of white matter, appearing dark grey in high resolution T2w and light grey in high resolution T1w (de Flores et al., 2020; Piskorowski & Chevaleyre., 2017). The challenges associated with imaging the hippocampus with sufficient quality for delineation are amplified in children, leading to limited availability of high-quality MRI suitable for hippocampal research. However, our work finds that the anatomical features of the hippocampus, such as the dark band in the T2w data, are equally visible in our child scans. The hippocampus exhibits high developmental variability during childhood and adolescence (Paus, 2005). Visibility of the SRLM is remarkable because myelination during development is argued to affect how the SRLM manifest in structural imaging (Milesi et al., 2014). Furthermore, paediatric hippocampal investigations are crucial given the implications of hippocampal development for memory research, and degeneration in clinical conditions such as epilepsy, and schizophrenia (Anahd & Dhikav., 2012).

Beyond the hippocampus our imaging protocol resulted in excellent fine structural contrast and visibility in the brain stem and the cerebellum. Our dataset overcomes barriers the delicate structure of the cerebellum presents, revealing the arbor vitae — the small branches of the cerebellum. Such advances open avenues for linking cerebellar structure to motor, cognitive, and clinical outcomes in both typical and atypical development. Further, anatomical differences have been identified in neurodevelopmental conditions including autism and ADHD (Stoodley, 2016). Efforts to characterise cerebellar organisation *in vivo* have long been hindered by resolution limits that obscure its intricate architecture (Mormina et al., 2017).

In stereotaxy, brain-based landmarks, particularly the AC and PC, are essential for orienting and localising neuroanatomical structures. The ACPC (intercommissural) line defines standard orthogonal planes in histology and serves as the reference for aligning MRI data to MNI space, enabling direct comparisons across modalities (Amunts et al., 2013; Conti et al., 2023; Mai & Majtanik, 2017). At conventional MRI resolutions (≥1 mm^3^), the PC’s distinctive “hook” morphology is not visible, leading to variability in its identification. In contrast, submillimetre imaging, such as ours, resolves the full hook, allowing for a more accurate comparison of data across subjects.

The satisfactory visibility of the PC introduces a novel challenge for future anatomical investigations: where exactly on or within the hook should the PC landmark be placed? For instance, a line drawn through the top of the hook versus the bottom differs in angle by only a few degrees at the PC. However, as the ACPC line drawn from these two points on the PC extends through the brain it yields systematically different coordinate systems and the error increases from a few degrees to many upon reaching the most lateral anatomy such as that of the cortex. Such variability undermines the core purpose of stereotaxic orientation, which is to provide a stable and transferable set of coordinates for anatomical reference, calling for an agreement to be made on the standardisation of location of the PC marker.

Subtle neuroanatomical differences associated with neurodevelopmental conditions often go undetected using conventional MRI resolutions. Achieving higher spatial resolution can improve measurement accuracy, reduce noise, and potentially lower the sample sizes required to detect significant differences. The methodological approach presented here demonstrates the feasibility of acquiring high-quality structural data in children. While our exemplars, the hippocampus, cerebellum, and PC, illustrate regions where such resolution is valuable, the broader utility of high-resolution MRI for studying the brain extends beyond. While we did not demonstrate the applicability of our methods to clinical populations, we suggest our methodology may also be suitable for some difficult-to-scan but cooperative populations.

Achieving 0.5 mm³ MRI images using conventional acquisition methods highlights the potential of 3T MRI. It may be possible to achieve even higher resolution at 3T through extended scan times and additional acquisitions. Given that access to high-performance computing is more readily available than higher-field MRI systems, optimising 3T protocols offers a practical alternative to deliver high-quality data typically requiring with ultra-high-field imaging. This increased resolution is central to visualising subcortical structures once limited to histological investigation, and although histology has much higher resolution at <0.01 mm³, MRI grows ever closer (Alkemade et al., 2023; Barazany & Assaf, 2012; Zarella et al., 2018). The ability to resolve intricate structures, such as the hippocampal subfields and cerebellar anatomy in paediatric neuroimaging, is closing the gap between *in vivo* MRI and *ex vivo* histology, providing a valuable tool for mapping brain development and studying its disruptions in various neurological and developmental disorders.

In conclusion we demonstrate the feasibility of acquiring sub-millimetre resolution images in paediatric populations, thereby extending the boundaries of what can be achieved *in vivo* during early development. The results demonstrate that careful optimisation of acquisition parameters and processing strategies can yield images of sufficient quality to delineate fine microstructural detail in the developing brain. All methods and processing workflows described in this study are openly available at https://osf.io/ckh5t/. By enabling access to such high-resolution paediatric neuroimaging, the present approach addresses long-standing technical barriers and provides a framework for both basic research and clinical translation.

## 5.1 Ethics

This study was approved by The University of New South Wales Human Research Ethics Committee (Reference: iRECS5939)

## 5.2 Data and Code Availability

All processing methods and scripts can be found at (OSF: https://osf.io/ckh5t/) and are compatible with Neurodesk which allowed for specification of software versions (Renton et al., 2024).

## 5.3 Author Contributions

BW: Conceptualisation, data acquisition, methodology, software and technical development, figure generation, formal analysis, manuscript preparation (draft + edit). MS: Conceptualisation, methodology, software and technical development, formal analysis, supervision, manuscript preparation (edit). GP: Funding acquisition, manuscript preparation (edit). SK: Conceptualisation, resources and infrastructure, supervision, manuscript preparation (edit).

